## Supplemental Figures and Tables for "Reprogramming feedback strength in gibberellin biosynthesis highlights conditional regulation by the circadian clock and carbon dioxide": 20250205_Supplemental Materials_GAHACR.docx

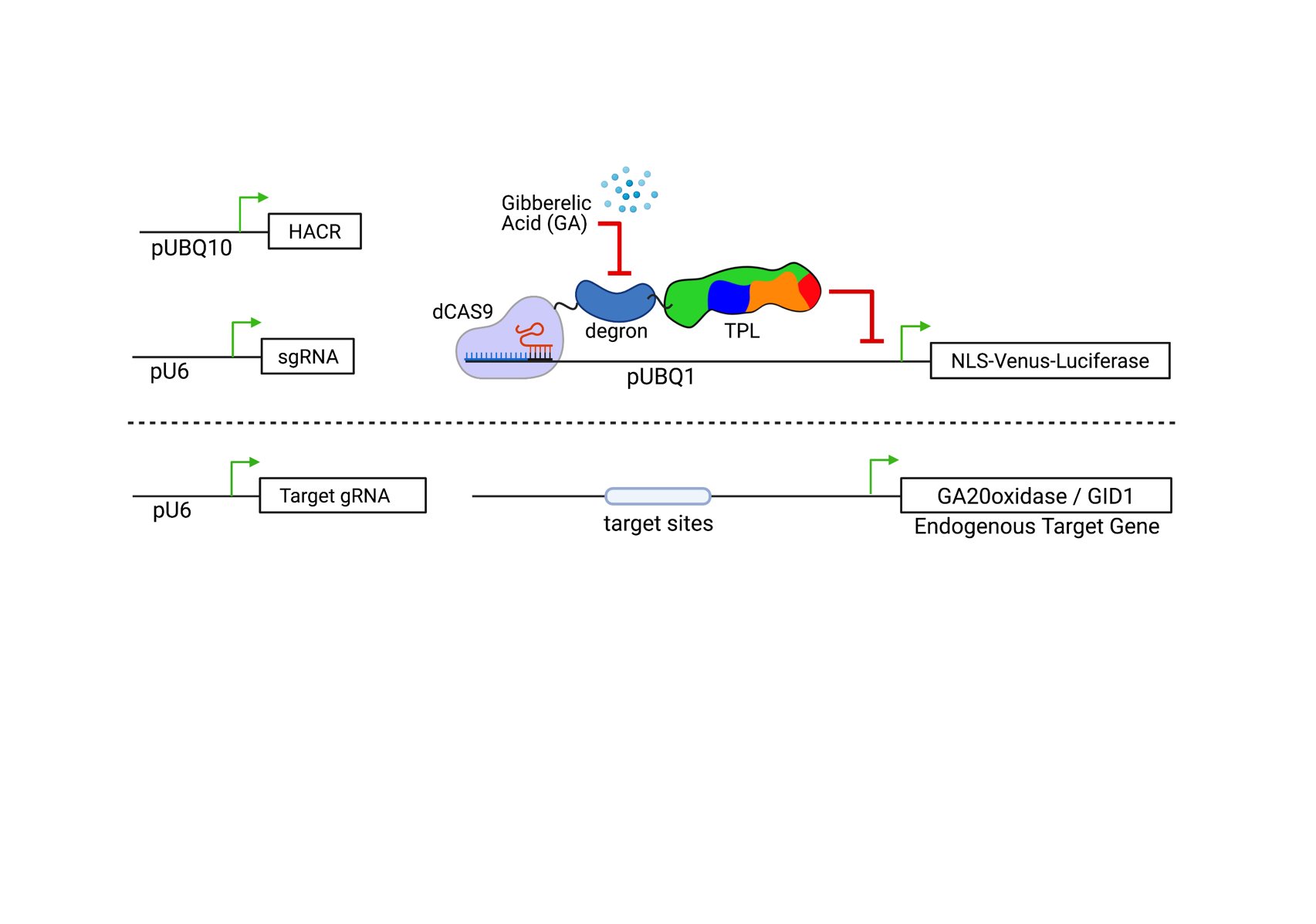

**Figure 2 Figure Supplement 1. Schematic of GA-HACR retargeting to endogenous genes in *Arabidopsis*.** The top schematic of the genetic circuit used to build GA responsive HACRs (above dotted line) was described in (Khakhar et al., 2018). In the lower portion of the schematic (below the dotted line) demonstrates the additional pAt-U6 driven gRNA which targets the endogenous genes (i.e. GA20ox or GID1). This allows the GA-HACR to simultaneously act as a reporter via Venus/Luciferase, but also modify existing genetic networks.

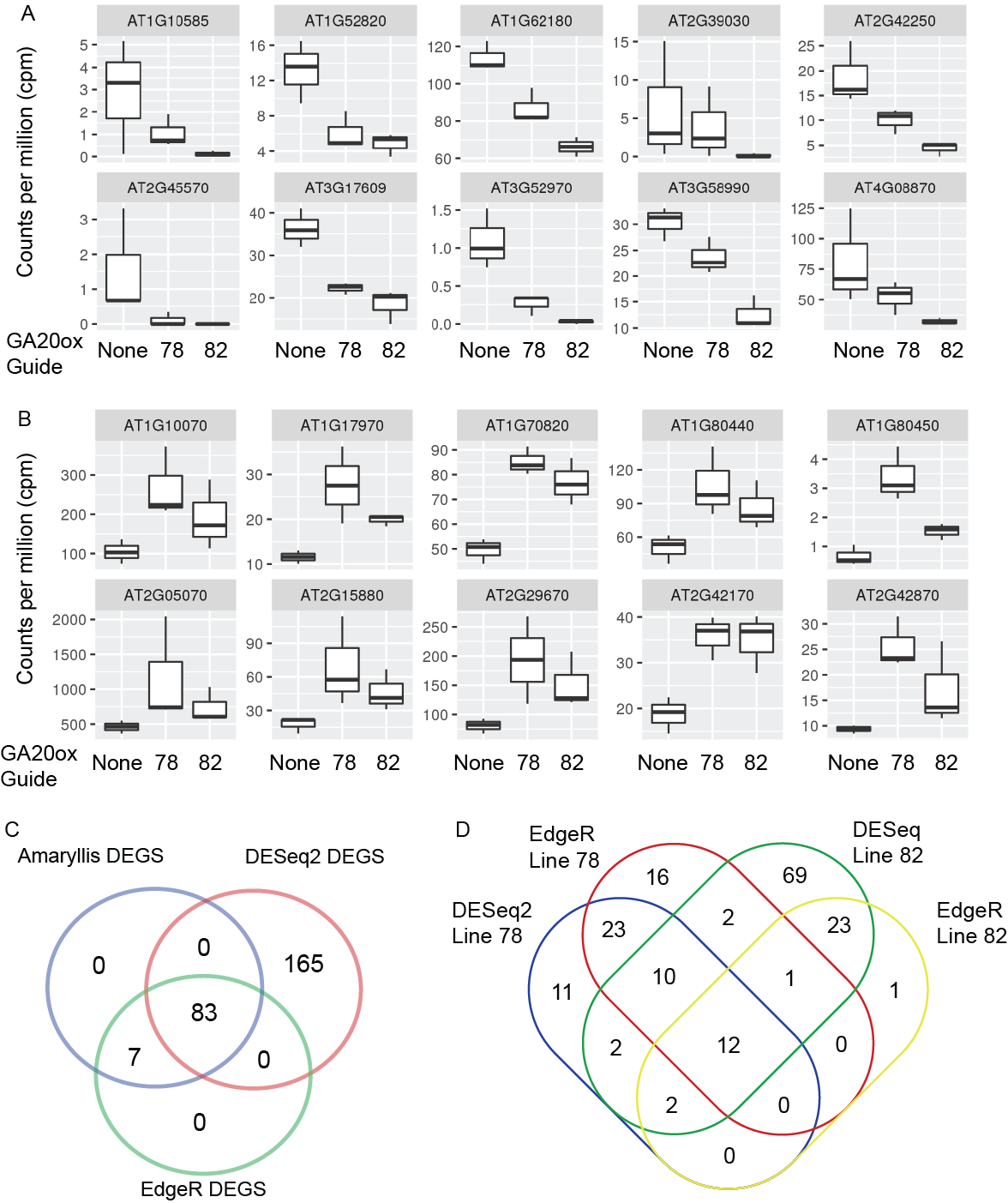

**Figure 3 Figure Supplement 1. RNA-Sequencing analysis of GAHACR network at ambient carbon dioxide levels – Identifying differentially expressed genes. A-B.** Selected DEGS were chosen to be graphed to demonstrate gene expression changes across the two selected lines (78 and 82) and wild type. We selected 10 down-regulated **(A)** and 10 up-regulated **(B)** DEGs, and graphed the counts per million across the 3 replicates in standard boxplots to demonstrate the similar trends across the two lines. **C.** To ensure that we were robustly detecting the maximum impact of the GA-HACR intervention, we applied two supplemental DEG finding packages, DESeq2 and EdgeR, and plotted the overlap in detected DEGs. **D.** A more detailed breakdown of the DEGs identified in the two DEG caller methods separated by genetic line.

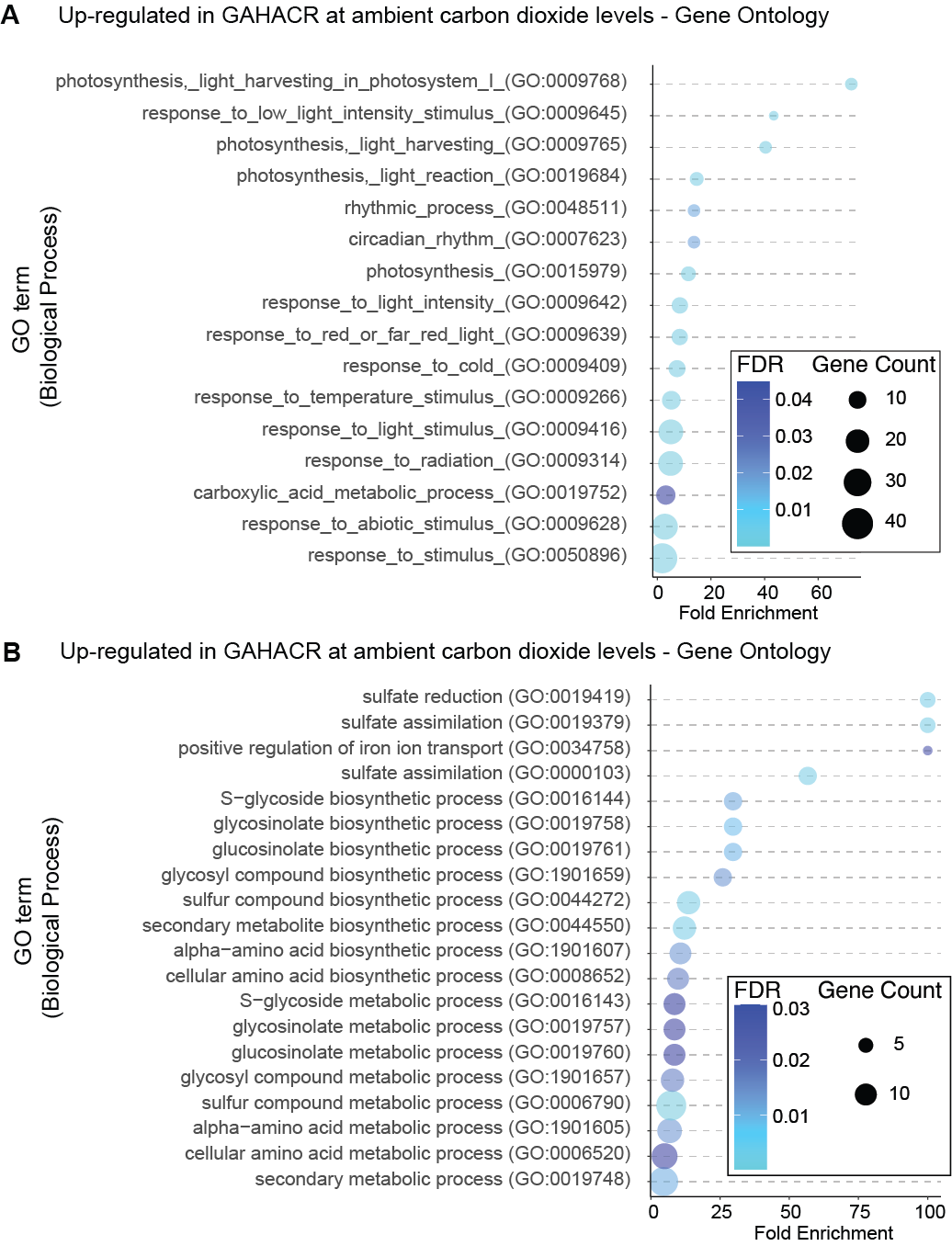

**Figure 3 Figure Supplement 2. RNA-Sequencing analysis of GAHACR network at ambient carbon dioxide levels – Gene ontology analysis.**

Gene Ontology terms enriched in the upregulated (A) and downregulated (B) differentially expressed genes from GAHACR targeted to *GA20ox* at ambient carbon dioxide levels. FDR – False discovery rate, GO was called via gProfiler. Data was graphed in R, using the ggplots2 package (see methods).

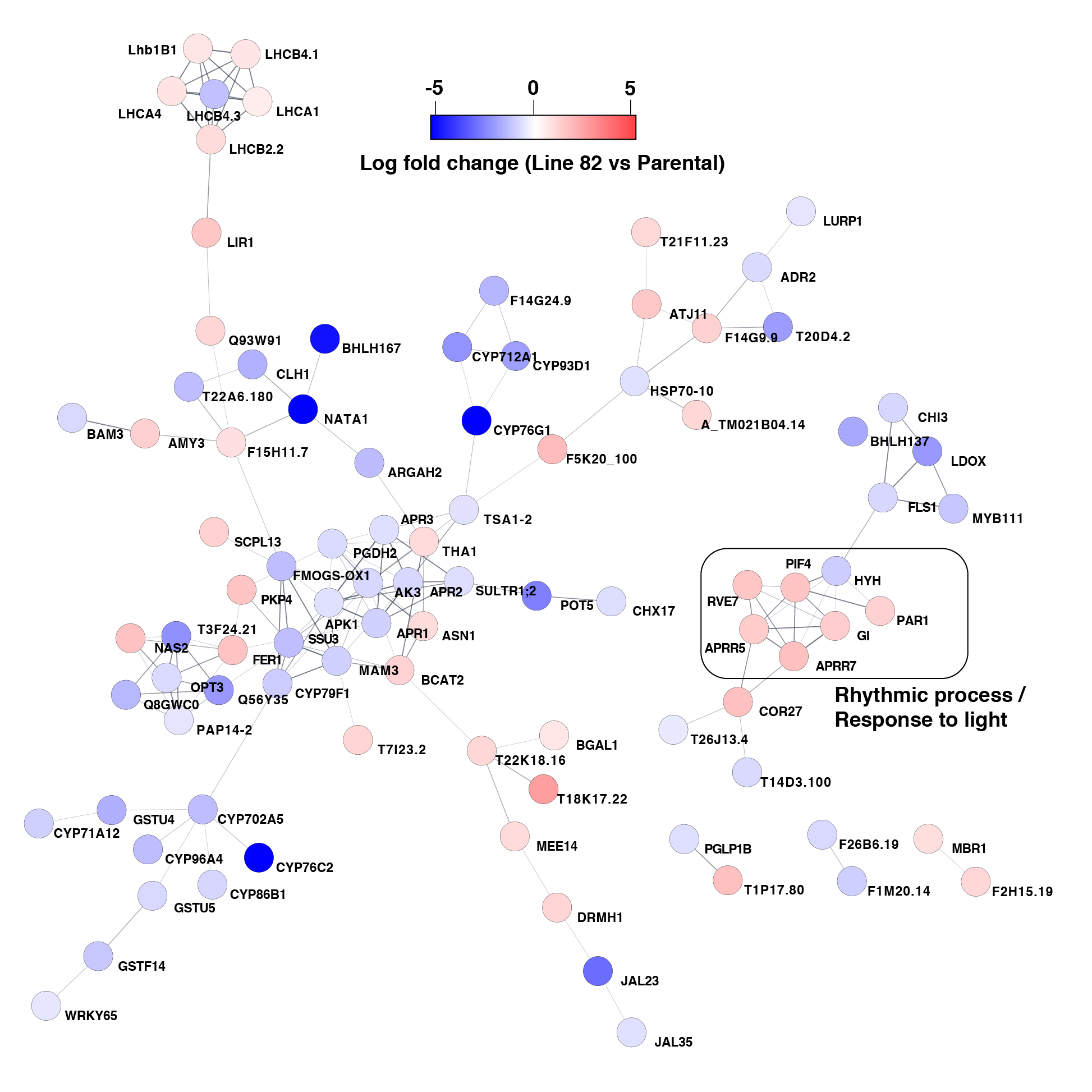

**Figure 3 Figure Supplement 3. Network analysis of the GAHACR network at ambient carbon dioxide levels.** DEGs from the GA-HACR RNA-sequencing performed at ambient carbon dioxide levels were imported into Cytoscape for network analysis using the STRING database. All singletons were trimmed, and functional enrichment was performed. A subnetwork of Rhythmic processes and Light responsive proteins were identified and excerpted into figure 3E. Nodes are color coded based on the log fold change observed in the GA-HACR lines 82 versus the parental GA-HACR line. DEGs that demonstrated a reduction in expression are colored blue and DEGs that increased expression are red (see scale).

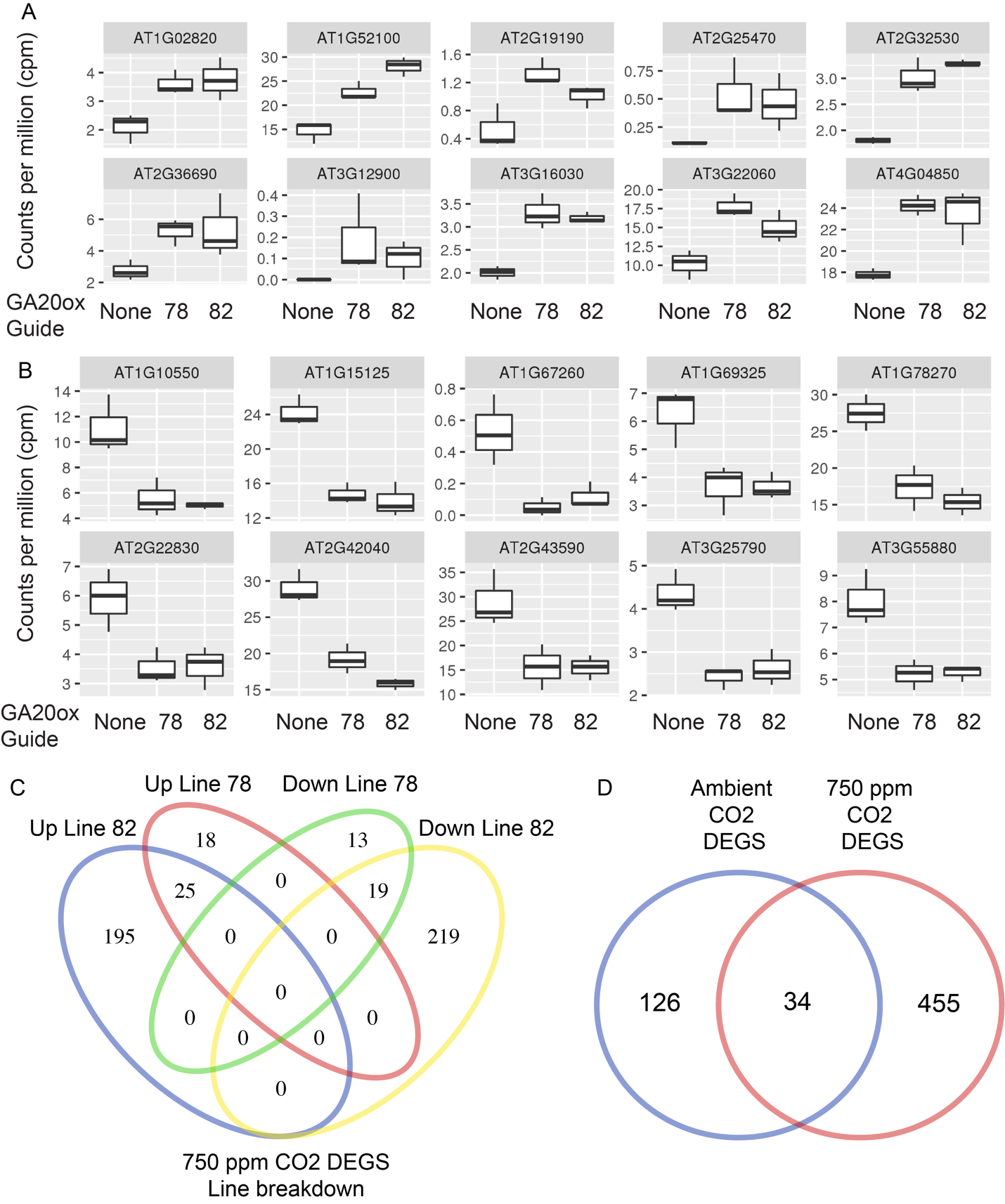

**Figure 3 Figure Supplement 4. RNA-Sequencing analysis of GAHACR network at elevated carbon dioxide levels – Identifying differentially expressed genes. A-B.** Selected DEGS were chosen to be graphed to demonstrate gene expression changes across the two selected lines (78 and 82) and wild type. We selected 10 down-regulated **(A)** and 10 up-regulated **(B)** DEGs and graphed the counts per million across the 3 replicates in standard boxplots to demonstrate the similar trends across the two lines. **C.** A more detailed breakdown of the DEGs identified in the two genetic lines at elevated carbon dioxide demonstrates the overlap in DEGs lists. **D.** The DEGS lists from the ambient and elevated (750ppm) carbon dioxide treatments were compared to determine if there is significant overlap between these datasets.

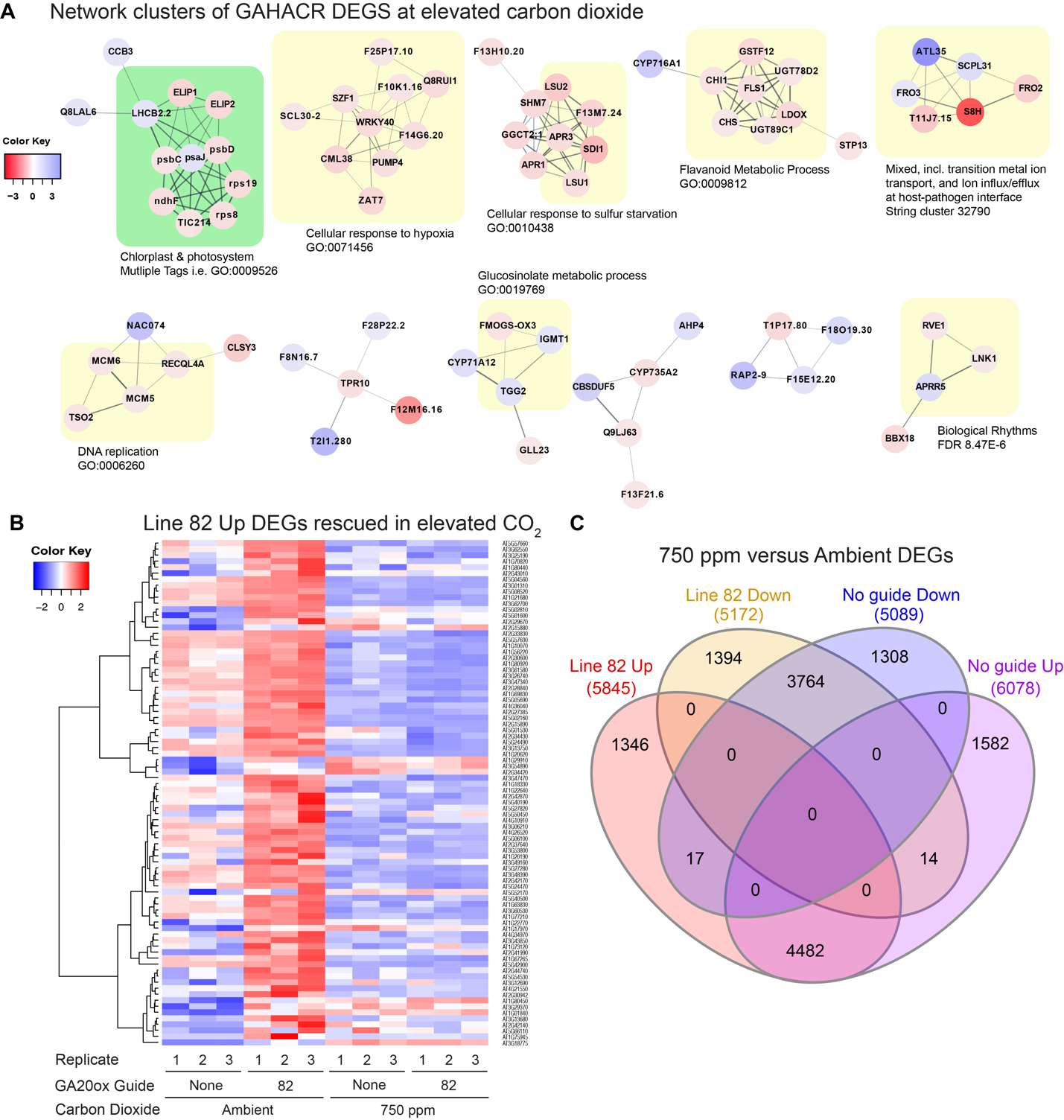

**Figure 3 Figure Supplement 5. Analysis of the GAHACR network at elevated carbon dioxide levels. A.** DEGs from the GA-HACR RNA-sequencing performed at elevated (750ppm CO_2_) carbon dioxide levels were imported into Cytoscape for network analysis using the STRING database. All singletons were trimmed, and functional enrichment was performed. The network was clustered using MCL (inflation value = 4). A subnetwork of Rhythmic processes was identified (bottom right, biological rhythms) that includes the genes PRR5 and RVE1. Nodes are color coded based on the log fold change observed in the GA-HACR lines 82 versus the parental GA-HACR line. DEGs that demonstrated a reduction in expression are colored blue and DEGs that increased expression are red (see scale). **B**. Heatmap of top upregulated DEGs identified by RNA-seq analysis at ambient CO_2_ levels (left 6 columns). Only line 82 upregulated DEGs are shown for simplicity, and many appear to be reduced in expression at elevated carbon dioxide levels (right 6 columns). Values are in Log2, where red is upregulated, and blue is downregulated. **C.** Intersection of DEGs generated by comparing ambient to elevated carbon dioxide demonstrate a large number of genes that are differentially expressed regardless of the GAHACR intervention.

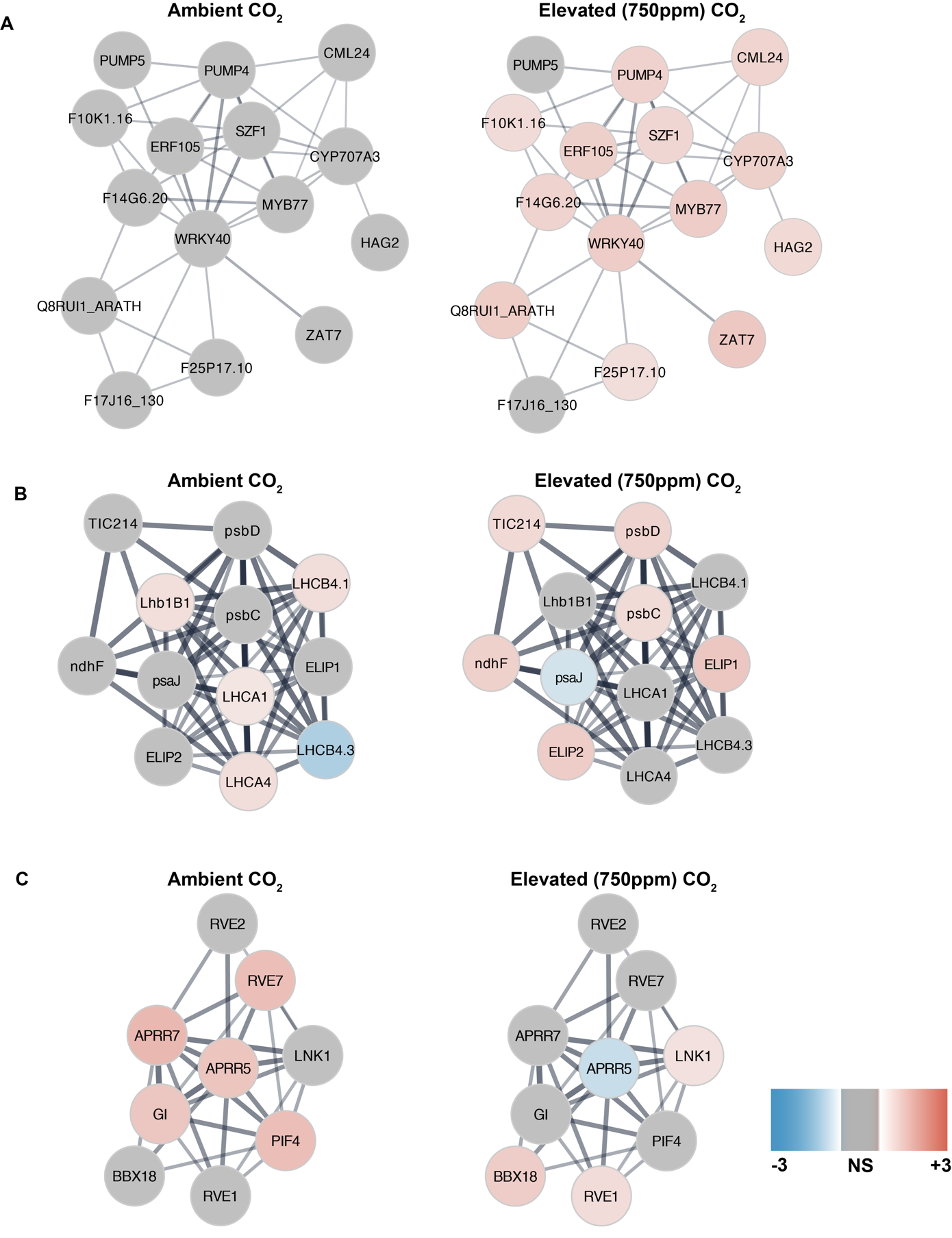

**Figure 3 Figure Supplement 6. Analysis of the GAHACR pooled network at both carbon dioxide levels.** DEGs from the GA-HACR RNA-sequencing performed at ambient and elevated (750ppm CO_2_) carbon dioxide levels were pooled and imported into Cytoscape for network analysis using the STRING database. All singletons were trimmed, and functional enrichment was performed. The network was clustered using MCL (inflation value = 4), and we manually subset selected clusters to examine how elevated carbon dioxide influences the networks. Each cluster node was colored based on the differential gene expression from the ambient (left column) or elevated (right column) no guide versus line 82 GAHACR experiments. **A.** Cluster defined by GO term Cellular response to hypoxia, GO:0071456, FDR = 1.02E-9. **B.** Cluster defined by GO Cellular component keyword Photosystem, GO:0009521, FDR = 3.12E-20 **C.** Cluster defined by UniProt keyword Biological Rhythms, KW-0090, FDR = 8.27E-16.
